# Persistent chromatin states, pervasive transcription, and shared *cis*-regulatory sequences have shaped the *C. elegans* genome

**DOI:** 10.1101/817130

**Authors:** James M. Bellush, Iestyn Whitehouse

## Abstract

Despite highly conserved chromatin states and *cis*-regulatory elements, studies of metazoan genomes reveal that gene organization and the strategies to control mRNA expression can vary widely among animal species. *C. elegans* gene regulation is often assumed to be similar to that of other model organisms, yet evidence suggests the existence of distinct molecular mechanisms to pattern the developmental transcriptome, including extensive post-transcriptional RNA control pathways, widespread splice leader (SL) trans-splicing of pre-mRNAs, and the organization of genes into operons. Here, we performed ChIP-seq for histone modifications in highly synchronized embryos cohorts representing three major developmental stages, with the goal of better characterizing whether the dynamic changes in embryonic mRNA expression are accompanied by changes to the chromatin state. We were surprised to find that thousands of promoters are persistently marked by active histone modifications, despite a fundamental restructuring of the transcriptome. We employed global run-on sequencing using a long-read nanopore format to map nascent RNA transcription across embryogenesis, finding that the invariant open chromatin regions are persistently transcribed by Pol II at all stages of embryo development, even though the mature mRNA is not produced. By annotating our nascent RNA sequencing reads into directional transcription units, we find extensive evidence of polycistronic RNA transcription genome-wide, suggesting that nearby genes in *C. elegans* are linked by shared transcriptional regulatory mechanisms. We present data indicating that the sharing of cis-regulatory sequences has constrained *C. elegans* gene positioning and likely explains the remarkable retention of syntenic gene pairs over long evolutionary timescales.

## Introduction

In the developing embryo, the differential control of RNA polymerase II (Pol II) transcription is crucial to establish major cell lineages, organize tissues, and shape morphological structures (Levine and Tjian 2003; Hamatani et al. 2004). Gene control mechanisms use *cis-*regulatory elements such as promoters and enhancers to integrate signals necessary for correct gene function (Levine et al. 2014). While minimal systems rely on competitive interactions between promoter-proximal DNA elements, transcription factors, and Pol II to regulate mRNA synthesis, eukaryotic genomes also utilize pre-mRNA splicing, chromatin structure, post-transcriptional RNA pathways, and long-range enhancer looping to modulate mRNA expression across different cell types and developmental stages (Banerji et al. 1981; Sharp 1985; Ptashne 2005; Cairns 2009).

Given the importance of promoters and enhancers in the context of cellular development and disease (Sur and Taipale 2016; Furlong and Levine 2018), recent high-throughput sequencing efforts have sought to annotate the genomic features of *cis*-regulatory regions across eukaryotes (Harbison et al. 2004; Gehrig et al. 2009; Negre et al. 2011; Shen et al. 2012; Daugherty et al. 2017). It is evident that certain “active” histone modifications like histone H3 lysine 4 methylation (H3K4me) and lysine 27 acetylation (H3K27ac) generally mark regions of chromatin accessibility and correlate with sites of transcriptional activity, allowing the approximate locations of most promoters and enhancers to be mapped (Heintzman et al. 2007). Histone modification landscapes generally reflect the transcriptional status of diverse cell types (Zhou et al. 2011)—this relationship is particularly evident during embryogenesis, which is marked by the progressive establishment and remodeling of the chromatin landscape as embryos become transcriptionally competent and cellular identities become defined (Lu et al. 2016; Hug et al. 2017).

Comparative transcriptome analyses of early metazoan development have revealed that, despite obvious morphological differences, the mRNA expression timing of orthologous genes is generally conserved across animal phyla (Domazet-Loso and Tautz 2010; Levin et al. 2016)– evidence suggests that common gene regulatory modules may have evolved to direct essential embryonic events, such as gastrulation, cell fate specification, and organogenesis (Lenhard et al. 2012; Boyle et al. 2014; Marletaz et al. 2018). Despite such similarity in mRNA patterns, it is clear that metazoan genomes employ vastly different structural and functional strategies to ensure that a given mRNA is present at the appropriate place and time (Romero et al. 2012). For example, genes in *Drosophila* are often regulated by interaction with transcriptional enhancers, which form stabilized DNA loops to insulate neighborhoods of differential Pol II transcription activity (Ghavi-Helm et al. 2014; Zabidi et al. 2015), whereas, trypanosomes appear to regulate gene expression post-transcriptionally and transcribe tens to hundreds of genes into a single polycistronic pre-mRNA (Clayton 2013). Genes co-transcribed into the same pre-mRNA are processed into individual mRNAs by splice leader (SL) trans-splicing, which couples 3’ polyadenylation of the upstream gene with the trans-splicing of a capped RNA leader to the 5’ end of a downstream gene (Jager et al. 2007). Examples of “multi-gene” transcription units have been documented in other organisms, such as *Euglena* (Moore and Russell 2012), *Arabidopsis* (Leader et al. 1997; Merchan et al. 2009), and mushroom forming fungi (Gordon et al. 2015). Based on these diverse transcriptional adaptations, it appears that the constraint of sharing *cis-*regulatory sequences by multiple genes in the same chromosomal space may fundamentally shape gene organization, Pol II transcription patterns, and post-transcriptional RNA control pathways (Koonin and Wolf 2010; Irimia et al. 2012; Silveira and Bilodeau 2018).

Because embryonic and post-embryonic cell lineages are largely invariant between individuals (Sulston and Horvitz 1977; Sulston et al. 1983), *C. elegans* has served as a powerful model system to characterize how gene expression states influence cell identity. *C. elegans* has developed multiple molecular mechanisms to optimize genome expression in response to developmental and environmental signals (Zaslaver et al. 2011; Maxwell et al. 2012; Warner et al. 2019), evidenced by the fact that *C. elegans* genes evolve at nearly twice the rate as their *Drosophila* orthologs (Mushegian et al. 1998; Coghlan 2005). Two notable features of *C. elegans* gene regulation are the organization of genes into operons and the extensive use of SL trans-splicing to process mRNA transcripts (Bektesh and Hirsh 1988; Blumenthal et al. 2002). Similar to the bacterial gene structure, *C. elegans* operons organize closely-spaced (<300 bp intergenic distance), colinear genes under the regulation of a single upstream promoter and a Pol II transcription start site (TSS)—however, the *C. elegans* genes contained in operons are not obligately co-expressed through translation (Zhao et al. 2004; Wang et al. 2010). Instead, polycistronic pre-mRNAs, (i.e. pre-mRNAs encoding multiple ORFs) are processed into individual mRNA messages by SL trans-splicing (Blumenthal et al. 2002). Transcriptome analyses established that 15% of *C. elegans* protein-coding genes, typically encoding growth functions and exhibiting germline-expression patterns, are organized into >1000 operons (Blumenthal et al. 2002; Reinke and Cutter 2009; Zaslaver et al. 2011)—however, considering that 70% of pre-mRNAs are SL trans-spliced (Zorio et al. 1994), it remains possible that additional *C. elegans* genes utilize polycistronic transcription and SL trans-splicing for expression (Morton and Blumenthal 2011). While the regulatory context for SL trans-splicing in *C. elegans* remains incomplete, similar gene regulatory adaptations have been identified in related nematode species, suggesting that common selective forces may have influenced developmental plasticity during nematode evolution (Kuwabara 1996; Webb et al. 2002; Stadler and Fire 2013).

Another striking distinction is the absence of CTCF or chromatin insulator proteins encoded by the *C. elegans* genome, suggesting that topological looping and higher order chromatin structures might be absent or fundamentally different (Heger et al. 2009; Dekker and Heard 2015). Genomes that encode chromatin insulators are capable of folding into compartments of differential Pol II transcription activity called topologically associating domains (TADs)—these compartments stabilize gene expression states by forming boundaries that prevent *cis-*regulatory elements from ectopically activating genes in neighboring TADs (Hnisz et al. 2016; Szabo et al. 2019). However, despite the lack of conserved TAD structures to regulate transcription, the chromatin landscape in *C. elegans* is decorated with the same patterns of patterns modifications found in other metazoa, including active marks: H3K27ac and H3K4me and repressive marks, H3K9me and H3K27me (Gerstein et al. 2010). Guided by the association between active histone marks and Pol II transcription initiation, recent chromatin profiling studies have revealed many thousands of putative gene promoters and enhancers across worm development (Chen et al. 2013; Daugherty et al. 2017; Ho et al. 2017). Yet, beyond basic characterization, we have a limited understanding of how chromatin may control the differential mRNA expression patterns during *C. elegans* development.

To examine the dynamics of Pol II transcription patterns and chromatin structure during *C. elegans* embryogenesis, we integrated maps of nascent RNA transcription, mRNA expression, and active chromatin modifications from highly synchronized embryo populations. By profiling embryos at distinct stages of development, we describe a highly dynamic transcriptional landscape that arises from a chromatin landscape that is largely invariant and may be inherited from the adult germline. Surprisingly, pervasive transcription of genes proximal to H3K27ac continues for the entirety of embryogenesis, even when mRNA levels are downregulated – a finding that highlights the extent to which *C. elegans* utilizes post-transcriptional mechanisms to control mRNA abundance. We also find that polycistronic transcription units represent a significant fraction of the embryonic transcriptome and cover many adjacent genes that are not part of operons. We propose that gene transcription is fundamentally regulated by *cis-*acting directional, elements that function to link adjacent genes. We present evidence that the gene regulatory systems employed by *C. elegans* have profoundly influenced gene organization.

## Results

### Chromatin landscape of early *C. elegans* embryogenesis

Although the genetic pathways and cellular events responsible for patterning the *C. elegans* embryo are well-documented (Sulston and Horvitz 1977; Sulston et al. 1983), a clear understanding of how the landscape of chromatin modifications and active Pol II transcription changes through early development is not known. Techniques sampling mRNA from whole embryos (Hashimshony et al. 2015; Boeck et al. 2016), individual tissues (Meissner et al. 2009; Tzur et al. 2018), and single-cells (Cao et al. 2017; Packer et al. 2019a) have revealed the dynamic expression of thousands of genes, but the gene regulatory context remains incomplete without integrating annotations of the chromatin architecture and Pol II transcription landscape at distinct developmental milestones (Gerstein et al. 2010; Levin et al. 2012; Kruesi et al. 2013; Ho et al. 2014; Pourkarimi et al. 2016; Daugherty et al. 2017).

In order to create chromatin state maps related to embryonic gene expression, we focused on H3K27ac and H3K4me2, as these modifications are widely considered to be indicators of RNA transcription at gene promoters and enhancers (Henriques et al. 2018). Embryo populations extracted conventionally by bleaching wildtype N2 hermaphrodites are developmentally asynchronous due to the staggered fertilization of oocytes in the adult germline (Tintori et al. 2016). To harvest developmentally synchronized populations of embryos, we utilized a hyperactive egg laying mutant, *egl-30(tg26)* (Bastiani et al. 2003; Pourkarimi et al. 2016)—as fewer embryos are contained in the uterus of gravid *egl-30(tg26)* adults, bleaching results in the isolation of pure populations of 1-4 cell embryos, compared to heterogeneous populations of early-gastrula stage (∼1-50 cells) embryos obtained from wildtype N2 adults (Bastiani et al. 2003). Since the purified embryos remain viable post-extraction, we developed embryo cohorts out to 60 min (“Early”), 200 minutes (“Gastrula”), and 600 minutes (“Late”) post-fertilization (PF) in order to assess histone marks of active transcription at distinct stages of morphogenesis (Supplemental Figure 1A). The three cohorts separate embryos into markedly different developmental stages: “Early” embryos display little transcription (Edgar et al. 1994) and are primarily composed of rapidly dividing cells; “Gastrula” embryos also contain dividing cells and a highly active transcriptome that supports rapid proliferation (Baugh et al. 2003); “Late” embryos are at the “three-fold” developmental stage and contain their near full complement of cells (∼550), as the transcriptome becomes devoted to expression of genes required for differentiation and organogenesis rather than proliferation (Chin-Sang and Chisholm 2000; Levin et al. 2012; Packer et al. 2019b). We validated the synchrony of our embryos by comparison of bulk population (n=60,000 embryos) mRNA with that derived from a single embryo time-course (Hashimshony et al. 2015), indicating that the embryonic transcriptome and developmental timing are recapitulated using this scaled-up approach (Supplemental Figure 1B).

### Stable maintenance of Pol II associated histone modifications throughout *C. elegans* embryo development

To construct transcription-associated chromatin state maps in early, gastrulation, and late stage *C. elegans* embryos, we prepared chromatin from synchronized cohorts of embryos through sonication and used standardized modEncode antibodies for immunoprecipitation (Methods) (Liu et al. 2011). As expected, both H3K27ac and H3K4me2 showed extensive overlap with each other and with intergenic promoter regions previously annotated by modEncode publications (Figure 1A) (Gerstein et al. 2010; Liu et al. 2011; Ho et al. 2014). One of the prominent but unexpected features revealed by our chromatin state maps is the highly similar pattern of H3K27ac and H3K4me2 modified nucleosomes across all embryo stages (Figure 1A). Typically, we find that *C. elegans* intergenic regions contain the same pattern of H3K27ac or H3K4me2 regardless of embryo stage (Figure 1A,B; Supplemental Figure 2A). The detection of transcriptionally-competent histone modification patterns in the earliest timepoints (60 min PF; 4-8 cells per embryo, Figure 1A) was puzzling as gene transcription is assumed to be relatively limited until gastrulation, when broad zygotic transcription initiates and the clearance of maternally-derived RNA transcripts is complete (Schauer and Wood 1990; Baugh et al. 2003; Levin et al. 2012).

**Figure 1:**
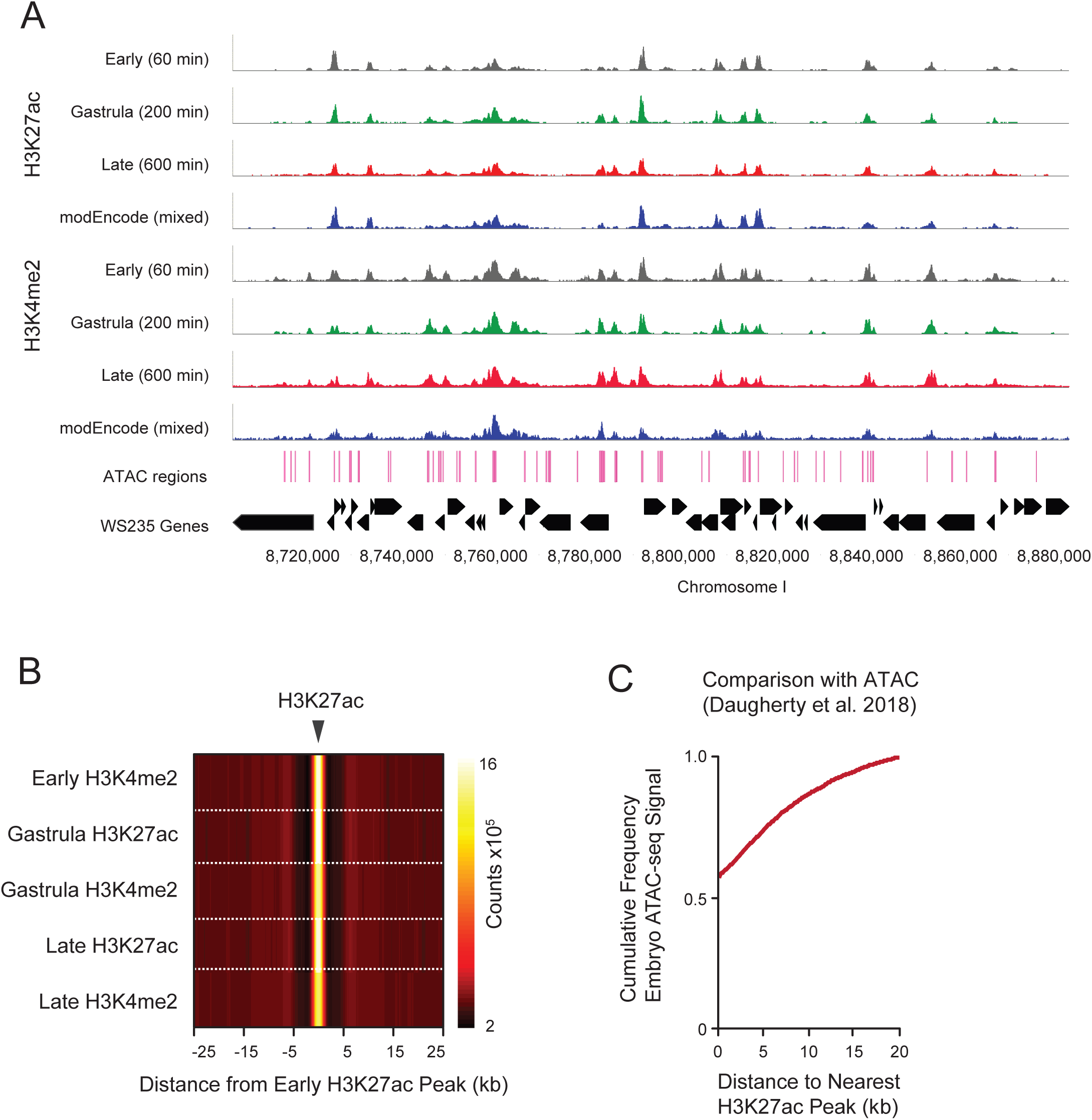
Chromatin domains of H3K27ac and H3K4me2 mapped during *C. elegans* embryogenesis are present soon after fertilization and are stably maintained into late stages of development. a) Synchronized populations of *C. elegans* embryos yield high-resolution maps of Pol II-associated chromatin marks. Populations of early embryos (black: 4-8 cells) were extracted from gravid *egl-30(tg26)* adults and developed *in vitro* in egg buffer to gastrula (green: 50-200 cells) and late embryonic (red: 200-550 cells) stages. Chromatin was extracted, sonicated, and immunoprecipitated by either H3K27ac or H3K4me2 antibodies prior to paired-end sequencing on Illumina Hi-Seq. ModEncode reference data from mixed N2 embryos (∼50-300 cells) is shown in blue, ATAC sites are shown as pink lines (Daugherty et al. 2017); WS235 gene annotations displayed in black. b) Active histone marks persist across embryonic stages. Normalized counts of ChIP-seq reads (red-yellow scale bar) from H3K27ac or H3K4me2 libraries were mapped in 100 bp genomic windows and graphed ±25 kb from early (4-8 cell) embryo H3K27ac midpoints. c) Synchronized ChIP-seq captures embryonic accessible chromatin states identified by ATAC-seq. Cumulative frequency plot of distances (up to 20 kb) between peaks of transposase accessible chromatin (Daugherty et al. 2017) and H3K27ac mapped in the early (4-8 cell) embryo; overlapping ATAC and ChIP-seq peaks are plotted at the origin (0 kb).

Since the chromatin landscape of early *C. elegans* embryogenesis has not been assayed extensively, we further characterized the genome-wide distribution of active chromatin marks. We sought to control for potential experimental artifacts caused by the sonication procedure by repeating the ChIP assay using chromatin sheared with micrococcal nuclease (MNase) and normalized read counts to a spike-in control (Orlando et al. 2014). The same reproducible H3K27ac and H3K4me2 patterns emerged by the MNase method, suggesting the open chromatin maps we generated are not a product of sonication artifacts (Supplemental Figure 2B). In addition, we considered the possibility that the stable H3K27ac and H3K4me2 ChIP-seq profiles generated from synchronized embryos may be disproportionately biased by the crosslinking and immunoprecipitation procedure (Teytelman et al. 2013). We compared embryo H3K27ac peak positions relative to accessible chromatin sites identified by a cross-linking independent assay, ATAC-seq (Daugherty et al. 2017). We found that the vast majority of transposase accessible sites identified in a mixed population of embryos to directly overlap (∼60%), or lie within 5 kb (∼70%) of our early embryo H3K27ac peaks (Figure 1C).

We annotated the embryo H3K27ac and H3K4me2 ChIP-seq profiles using the MACS (V2.1) peak calling algorithm to examine how early chromatin marks become distributed once zygotic transcription is activated during gastrulation and morphogenesis (Zhang et al. 2008). By plotting the normalized counts of H3K27ac and H3K4me2 signal from each embryonic time point relative to H3K27ac peaks annotated in the 4-8 cell embryo, we found these histone modifications lie within highly consistent genomic intervals from early embryogenesis (60 min PF) through the late stages of morphogenesis (600 min PF) (Figure 1A, 1B). Moreover, we find that other transcription-associated chromatin marks detected in L1 and L3 stage worms, including H3K4me3, H3K18ac, and H3K23ac, were also highly enriched within the same genomic intervals as persistent, early embryo H3K27ac marks (Supplemental Figure 2C).

While vertebrate embryogenesis requires extensive erasing and rewriting of active histone marks during developmental transitions (Dahl et al. 2016; Zheng et al. 2016; Zhang et al. 2018), our chromatin maps and enrichment analysis suggest that the histone modification status of early embryonic promoters (4-8 cell, pre-gastrula stage) appear to be retained, not erased, in somatic tissues of differentiated embryos. Despite massive changes in mRNA expression programs during *C. elegans* development (Levin et al. 2012; Hashimshony et al. 2015; Packer et al. 2019a), a large fraction of accessible chromatin domains in the early embryo are maintained throughout late embryo morphogenesis and even into post-embryonic development (Supp. Figure 2C).

**Figure 2:**
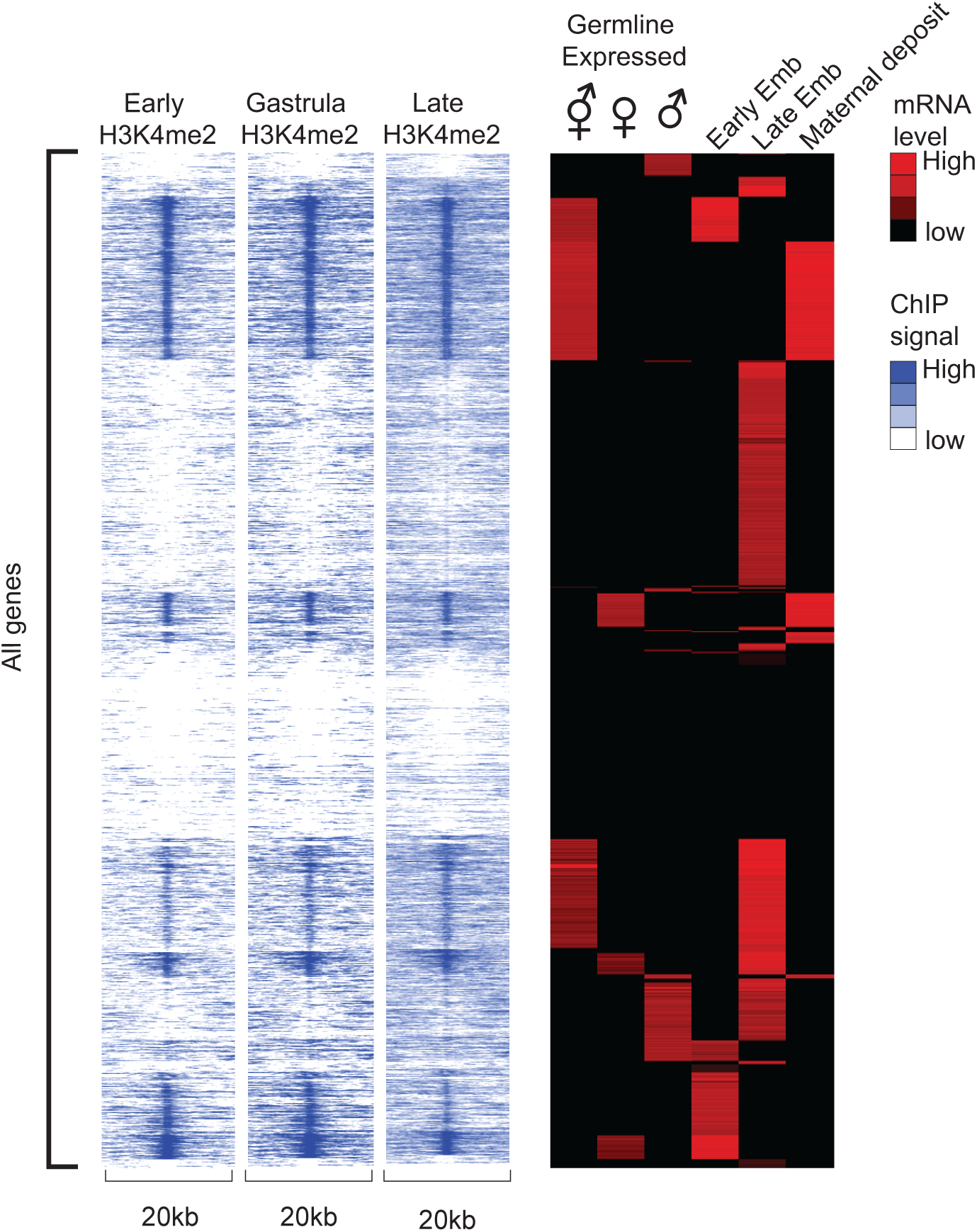
Association of embryonic histone modifications with germline expressed genes mRNA abundance was calculated for genes expressed in the germline (Ortiz et al. 2014) or embryo (Hashimony et al. 2015). Germline data was segregated according to mRNA enrichment in female (♀) male (♂) or both (♂♀) germ cells. Embryonic data was segregated into transcriptomes predominantly representing maternally deposited, early embryo (expressed 100-300 minutes), or late embryo (expressed 600-800 minutes) profiles. Data was clustered into 20 nodes according the maximal abundance stage and abundance; resulting gene order is displayed as a heatmap on right. For each gene the abundance of histone modifications at three stages of development was calculated for a 20kb range centered on the annotated start site for the first exon. The abundance of histone modifications (H3K4me2) is displayed as heatmaps that maintain gene order; analysis with H3K27ac nearly identical.

### Germline transcription may template modifications in the early embryo

We had anticipated that histone modifications of Pol II transcription would appear progressively during embryogenesis, in a manner similar to the activation of *cis-*regulatory regions mapped in *Drosophila* and mammalian embryos (Blythe and Wieschaus 2016; Hug et al. 2017; Wu et al. 2018). However, we find evidence that H3K27ac and H3K4me2 are present in the earliest stages of embryogenesis (60 minutes PF) prior to the broad phase of transcriptional upregulation at gastrulation. Multiple lines of evidence suggest that *C. elegans* embryos can retain parental histone modification patterns (including H3 acetylation and H3 lysine 4 methylation) during the period spanning fertilization and early embryonic cell divisions (Arico et al. 2011; Li and Kelly 2011; Kelly 2014). The maintenance of chromatin accessibility in the embryo may be related to transgenerational epigenetic inheritance, the phenomenon by which gene regulatory information is passaged through the adult germline to the fertilized zygote (Rechtsteiner et al. 2010; Li and Kelly 2011; Gassmann et al. 2012). It is estimated that well over 50% of the *C. elegans* genome is expressed at some stage by mitotic or meiotic germ cells, while a substantial proportion of RNA transcripts detected in the syncytial germline have been shown to be subsequently expressed zygotically during embryogenesis (Baugh et al. 2003; Reinke et al. 2004; Ortiz et al. 2014; Hashimshony et al. 2015; Tzur et al. 2018). Because overlapping expression patterns can complicate assignments of a germline or zygotic origin for early embryonic RNA transcripts, we asked whether specific germline-gene expression patterns may have influenced the accessible chromatin state in the *C. elegans* embryo.

*C. elegans* hermaphrodites produce oocytes and sperm within the same syncytial gonad, however well-characterized mutants in the sex-determination pathways have been used to differentiate oogenic [(*fog-2* (*q71*)], spermatogenic [(*fem-3 (q96)*], and gender-neutral transcriptomes (not enriched in either mutant) in the hermaphrodite germline (Ortiz et al. 2014). When we plotted the distance between germline-expressed genes and the nearest upstream H3K27ac peak in the early embryo, we found a notable enrichment of oogenic and gender-neutral genes downstream of early embryo active chromatin (Supplemental Figure 3), supporting the model that histone marks linked to Pol II transcription of genes in the adult germline can be retained in pre-gastrula embryos (Rechtsteiner et al. 2010; Arico et al. 2011; Kishimoto et al. 2017). In contrast, spermatogenic gene promoters show the weakest association with active histone marks in the early embryo, an outcome supported by evidence that transcription-associated histone acetylation and methylation on *C. elegans* sperm genes is erased prior to the onset of zygotic transcription in the embryo (Katz et al. 2009; Samson et al. 2014; Tabuchi et al. 2018). Given the close association between germline expressed genes and histone modifications found in the embryo, we next surveyed how embryonic histone modifications change in relation to the time at which an mRNA for a particular gene is abundant. For this analysis, we defined whether a gene is expressed in the germline (Ortiz et al. 2014) and whether that gene mRNA is also maternally deposited, expressed in early, or expressed in late embryogenesis (Hashimshony et al. 2015). We then clustered genes according to their mRNA abundance at various germ and/or embryonic cell stages and mapped the abundance of active histone marks directly upstream of individual genes in each cluster. As shown in Figure 2, this analysis reveals that many germline expressed genes (except for spermatogenic genes) appear to retain histone modifications into the embryo—we find the enrichment of active marks is the strongest at germline gene promoters that are also embryonically transcribed, yet it is notable that the *C. elegans* accessible chromatin state at many gene promoters persists regardless of mRNA expression time.

### Broadly-expressed and developmentally-regulated *C. elegans* genes are housed in distinct open chromatin states

Studies linking mRNA expression patterns to cellular events of metazoan embryogenesis have described two major developmental transcriptome phases: an early, proliferative stage and a late, differentiating phase (Levin et al. 2016). To assess the spatial and temporal mRNA expression patterns at differentially-marked chromatin regions during embryogenesis, we grouped modification peaks based on their stability across embryonic stages. Specifically, we classified two developmental contexts for embryonic H3K27ac and H3K4me2 domains: stable active chromatin (SAC), describing accessible chromatin regions (H3K27ac: n=3387, H3K4me2: n=5168) that are fixed in position and signal from gastrulation (200 min PF) to morphogenesis (600 min PF) (Supp. Table 1), and dynamic active chromatin (DAC), which defines new sites of H3K27ac (n= 1971) or H3K4me2 (n=2009) established only in the late embryo (Figure 3A, Supp. Table 2). We focused on H3K27ac (H3K4me2 gives near identical results, not shown) and assigned each DAC or SAC peak to its closest *C. elegans* gene and plotted the mRNA log_2_ fold change value of each respective gene from gastrulation to late embryogenesis (Hashimshony et al. 2015).

**Figure 3:**
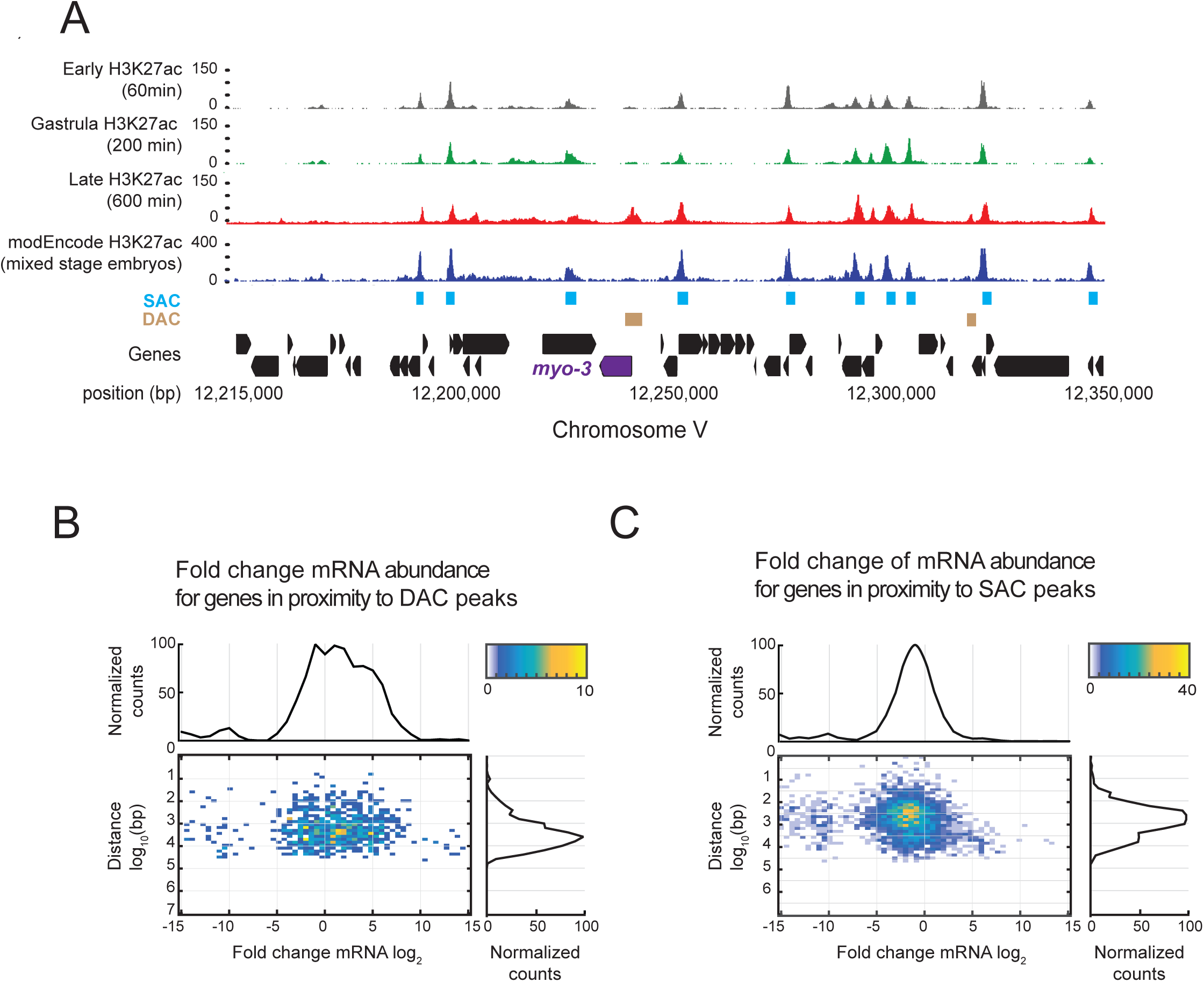
Appearance on new sites of histone modification are correlated with mRNA expression changes during *C. elegans* embryogenesis a) Histone modifications in *C. elegans* embryo were classified into stable active chromatin (SAC) or dynamic active chromatin (DAC) regions. Early (black), gastrula (green), and late (red) embryo H3K27ac tracks are displayed along with annotated features—SAC domains (blue boxes), defined as peaks which persist between gastrulation and late embryogenesis (overlap = 0.5), and DAC domains (tan boxes) that appear in late embryos. The *myo-3* gene which is expressed late in embryogenesis is highlighted in purple. WS235 annotated genes are displayed in black b) DAC peaks associated with mRNAs expressed late during embryo morphogenesis. mRNA log_2_ fold change values (x-axis), representing changes in gene mRNA counts between 200 min and 600 minutes, were calculated from the Hashimony et al. 2015 dataset and plotted on the heatmap at the gene’s respective distance from the nearest DAC peak (n=1971) (y-axis). Color scale indicates number of genes at that coordinate. C) SAC peaks marking stable H3K27ac domains represent sites of mRNA downregulation during transition to morphogenesis. mRNA fold change values were calculated as described previously and plotted with respect to gene distance from the nearest SAC peak (n= 3887). Color scale indicates number of genes at that coordinate.

The analysis highlights strikingly different mRNA expression profiles associated with DAC (late-appearing) and SAC (stably-maintained) chromatin regions during embryogenesis. Figure 3B reveals that genes closest to DAC sites are mostly upregulated in the late embryo (positive log_2_ fold change) and exhibit cell-type specific expression patterns— gene ontology analysis indicates that DAC-associated genes are repressed in most cells during early embryogenesis and activated in differentiated cell types (e.g. neuron, muscle) during morphogenesis (Supplemental Table 3a) (Perez-Lluch et al. 2015; Evans et al. 2016; Levin et al. 2016). Contrasting with the DAC regions, genes directly downstream of SAC regions experience a clear decrease in mRNA abundance in the late embryo (negative log_2_ fold change) (Figure 3C)—SAC-associated genes exhibit broad tissue expression patterns and encode cellular housekeeping functions such as growth, metabolism, and cell proliferation (Supplemental Table 3b), explaining the steep drop in mRNA levels in late embryos which have ceased cell divisions.

### Pervasive transcription of unstable, growth mRNAs in the differentiated late embryo

Given that polyadenylated mRNA is an insufficient proxy for the true extent of genome transcription, we tested the possibility that sites of persistent H3K27ac histone modification remained transcriptionally active through embryogenesis. Thus, we mapped the distribution of transcriptionally engaged Pol II genome-wide, using the global run-on sequencing (Gro-seq) technique using nuclei from synchronized populations of early and late C. elegans embryos (Core et al. 2008; Kruesi et al. 2013; Core et al. 2014) (Materials and Methods). We made several alterations to our Gro-seq approach to ensure that sequenced reads accurately represent the nascent coding and non-coding transcriptome of *C. elegans* embryogenesis. As most of the BrUTP will be incorporated into ribosomal RNA transcription, we depleted ribosomal RNA (rRNA) from affinity purified nascent RNA transcripts prior to cDNA synthesis. We also enriched for transcripts retaining their nascent 5’ end by not fragmenting our immunopurified RNA prior to cDNA synthesis— this step was necessary in previous Gro-seq approaches designed for the Illumina HiSeq short-read sequencing platform (Kruesi et al. 2013). We prioritized the selection of full-length nascent RNA transcripts by using a highly processive reverse transcriptase to synthesize cDNA libraries, which was subsequently size selected for transcripts >300 bp (Methods). Long cDNAs were sequenced using the MinION cDNA-sequencing platform from Oxford Nanopore Technologies—longer cDNA sequencing reads afforded greater coverage of the genome to identify gene regulation by RNA polymerase, but also minimized background signal common to PCR amplified short-read sequencing libraries.

To analyze the pattern of nascent RNA transcription throughout embryogenesis, we sequenced 3.53 GB and 5.32 GB of cDNA from gastrulation and late stage embryo nuclei, respectively. Less than 2 % of our Gro-seq reads include *C. elegans* splice leader sequences SL1 or SL2 (Allen et al. 2011), suggesting that the enrichment of nascent BrU-labeled RNA and a size selection for longer transcripts revealed a transcriptome before trans-splicing of outrons and monocistronic mRNA processing, including 3’ end cleavage and polyadenylation (Evans et al. 2001; Saito et al. 2013; Garrido-Lecca et al. 2016). We find much of our nascent RNA signal to be overlapping or immediately downstream of active histone marks, a positioning we would expect for transcriptionally engaged RNA polymerase (Figure 4A) (Core et al. 2014). Supporting these observations, we also found that Pol II TSSs from embryos, mapped by Gro-cap sequencing (Kruesi et al. 2013), overlap both SAC and DAC chromatin states (Figure 4B), highlighting that the active chromatin regions we have mapped are reliable indicators of Pol II TSSs. Although our Gro-seq format sequenced full-length nascent RNA transcripts, we observed a high degree of similarity to the Kruesi et al. profile of embryonic Pol II transcription, in terms of transcript density at promoter regions, coverage of gene bodies, and downstream intergenic transcription (data not shown).

**Figure 4:**
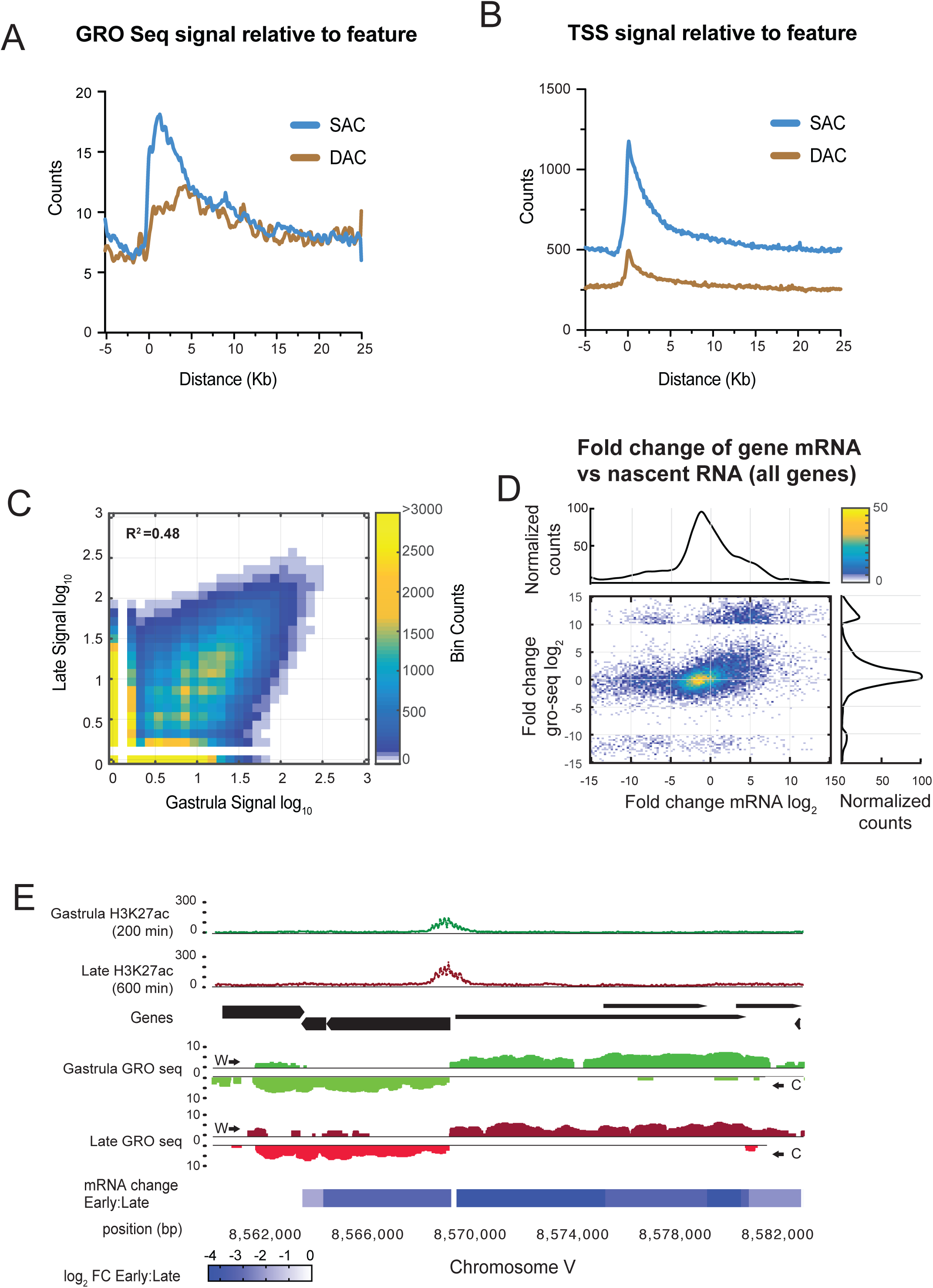
Nascent RNA sequencing reveals chromatin regulatory landscape for Pol II transcription and mRNA stability during *C. elegans* embryogenesis a) Pol II transcription downstream of SAC and DAC domains. Strand specific counts from mapped late embryo Gro-seq libraries were binned in 100 bp genomic windows with respect to the midpoint of the nearest SAC (blue) or DAC (tan) peak. b) As A, except data from Kruesi et al. 2013, which identified TSSs by mapping 5’ capped nascent RNA was plotted relative to SAC (blue) and DAC (tan) midpoints. c) Pol II transcription patterns highly consistent between early and late embryogenesis. Genome was broken into 100bp bins and signal for each bin was plotted in gastrula and late embryos. d) Major transcriptome remodeling during *C. elegans* embryogenesis involve drastic changes in mRNA stability, yet persistent nascent Pol II transcription patterns. The log_2_ fold change for mRNA and nascent RNA was calculated for each gene and then plotted. Color scale indicates number of genes at that coordinate. e) Genome browser image that highlights change in mRNA is not necessarily a result in decrease of transcription. Top, histone modification patters captured from gastrula and late embryos. Middle, nascent RNA mapped by Gro-seq is shown for gastrula (green) and late (red) embryos. Reads mapping to the Watson or Crick strands are shown above and below the center lines, respectively. Bottom, heatmap to represent the log_2_ fold change in mRNA abundance at genomic locus after transition from gastrula to late stage embryos.

Comparison of Gro-seq profiles from the gastrula and late embryos revealed consistent patterns of nascent RNA transcription across the genome (Figure 4C), which is in keeping with the persistent patterns of active histone modifications described above. We then examined the relationship between nascent and polyadenylated mRNA transcripts by directly comparing the fold change of mRNA against fold change in nascent RNA at all genes during the transition from early to late embryogenesis. Despite global changes in mRNA abundance, Figure 4D reveals that many genes experience little change in Pol II transcription levels— with notable exceptions: 1) genes newly transcribed by Pol II in late embryos (positive FC in mRNA and nascent RNA: upper-right quadrant) and 2) genes which completely lose Pol II transcription (negative FC in mRNA and nascent RNA: lower-left quadrant. The remaining genes exhibit a far different regulatory pattern characterized by persistent nascent RNA transcription between early and late embryos, but a major downregulation of polyadenylated mRNA levels, suggesting an extensive utilization of post-transcriptional mechanisms to control mRNA abundance during the transition from early to late embryogenesis (Figure 4E).

### Differential mRNA expression from polycistronic transcription units

Observation of Gro-seq patterns at consistently transcribed SAC regions revealed a tendency for neighboring, colinear genes to be transcribed continuously in the same transcription unit (Figure 5A). The *C. elegans* genome is rare among animals given the tendency for genes to be arranged into operons, which transcribe multiple open reading frames (ORFs) into a single polycistronic transcript (Spieth et al. 1993; Zorio et al. 1994; Allen et al. 2011). Individual mRNAs within operons are processed from polycistronic pre-mRNAs by splice leader (SL) trans-splicing, which mechanistically couples the 3’ end cleavage of the upstream mRNA to the 5’ capping of downstream mRNAs (Morton and Blumenthal 2011; Garrido-Lecca et al. 2016). An estimated 70% of RNA transcripts are thought to be processed by SL trans-splicing and might be of polycistronic origin, however the genomic regulatory context in terms of *C. elegans* development remains unclear. We decided to map continuous stretches of Pol II transcription (transcription units) genome-wide in order to examine how transcriptional and post-transcriptional RNA regulatory patterns may have influenced embryonic gene expression patterns (Merritt et al. 2008; Elewa et al. 2015; Lima et al. 2017; West et al. 2018).

**Figure 5:**
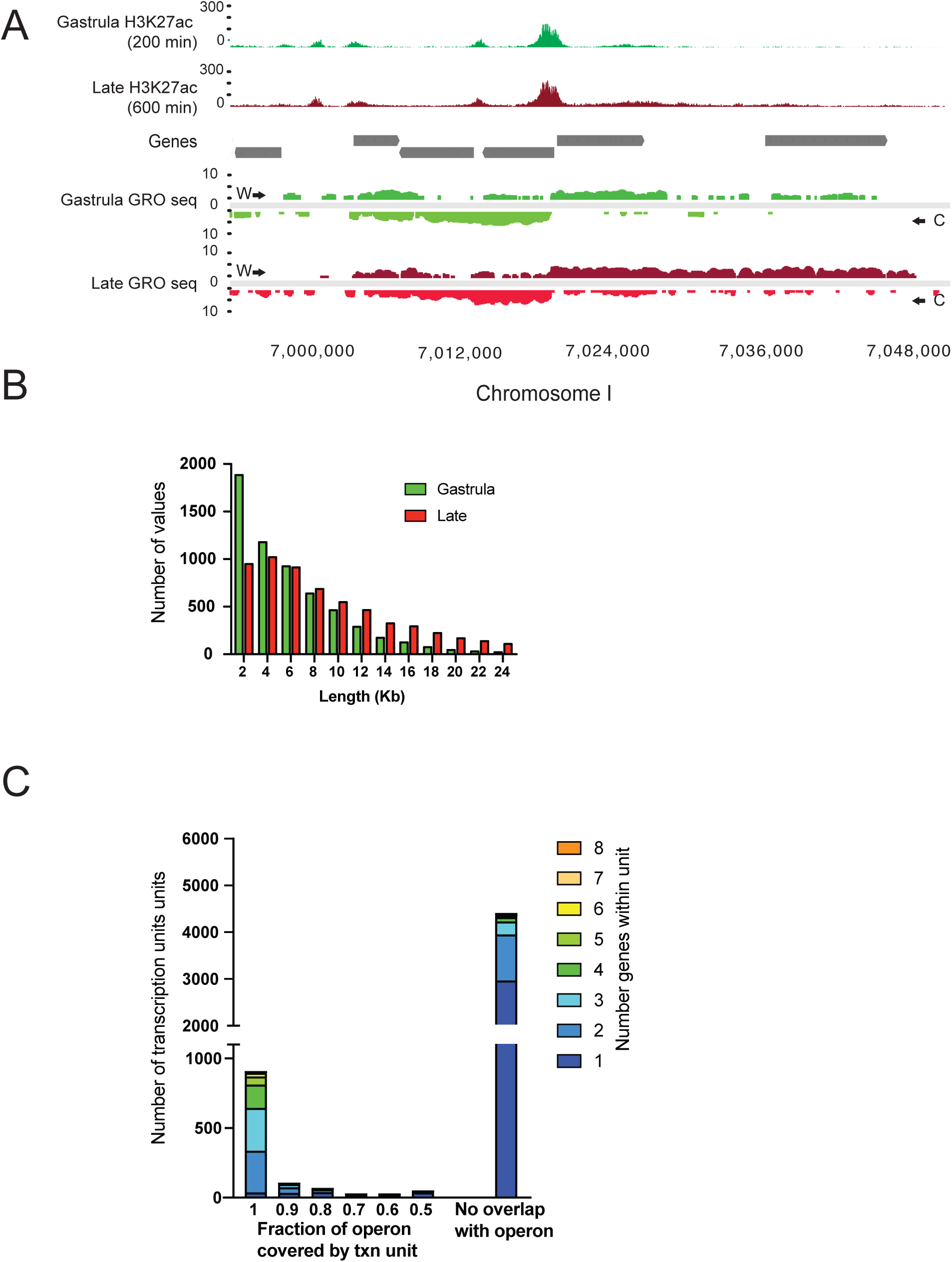
Identification of polycistronic RNA transcription. a) Genome browser image that highlights the extent of nascent RNA transcription during embryogenesis. Top, histone modification patterns captured from gastrula and late embryos. Middle, WS235 gene coorindates. Bottom, nascent RNA mapped by Gro-seq is shown for gastrula (green) and late (red) embryos. Reads mapping to the Watson or Crick strands are shown above and below the center lines, respectively. b) Mapped transcription units increase in length as development proceeds. The lengths of all transcription units was determined and the frequency of those lengths plotted. c) Chart showing the prevalence of operons in polycistronic transcription units. The gene content of each transcription unit mapped in the late embryo was analyzed to reveal the fraction of genes that are contained within an operon. Despite being associate with growth, most operons have all genes transcribed. Several thousand transcription units do not contain genes defined as part of operons i.e. 0 coverage. These transcription units primarily cover single genes, but ∼1500 span 2 or more coding sequences.

We utilized gro-HMM, which uses a two-state hidden Markov model (HMM) to annotate the boundaries of transcribed and non-transcribed units genome-wide (Chae et al. 2015). We sought to minimize the merging of unrelated transcriptional noise into our annotations of protein coding gene transcription—therefore, we fit a model which minimized error in background transcript calls and split up transcription unit mergers which were not supported by Gro-seq evidence. Furthermore, we adjusted model parameters to account for *C. elegans* genome size and gene density, which corrects gro-HMM calculations of Pol II transition from transcribed to non-transcribed states (Chae et al. 2015).

The majority of the gro-HMM defined transcription units overlap protein coding sequences: 69% (n= 6643) of the early embryo and 53% (n=6573) of the late embryo (Supplemental Table 4). The median length of transcription units increases in the transition from early to late embryogenesis, supporting the general observation of increased Pol II transcription in late embryos (Figure 5B). While most studies of *C. elegans* polycistronic transcription are limited to previously annotated operons (<300 bp intergenic distance) (Zorio et al. 1994; Blumenthal et al. 2002), unusual transcription units covering multiple, neighboring genes separated by kilobases of intergenic space have also been described (Morton and Blumenthal 2011). Using the gro-HMM annotations, we could identify that the majority of operons are encompassed by nascent Pol II transcription units (Figure 5C: operon fraction=1, (Allen et al. 2011)). Furthermore, annotation of late embryo Gro-seq profiles identified more than 1500 transcription units that encompass multiple genes that are not included in known *C. elegans* operons. While it is challenging to define whether or not transcription that covers multiple genes is composed of multiple overlapping transcription units, the extent to which the intergenic regions separating coding sequences is transcribed is notable, as is the frequent lack of detectable active chromatin in the transcribed intergenic regions.

### Conservation of colinear arrangement of genes in transcription units

Genes are not randomly positioned across genomes (Nguyen and Bosco 2015). Despite being separated by large evolutionary distances, animal genomes typically retain large blocks of orthologous genes whose composition and order has been conserved (Liu et al. 2018). Such regions of synteny are often bounded by cis-regulatory sequences; current models suggest that genes in syntenic regions may share transcriptional regulatory elements that ensure gene positions can become linked in transcriptional space and constrained by nearby expression patterns (Kikuta et al. 2007; Perry et al. 2010; Birnbaum et al. 2012).

We reasoned that if *C. elegans* transcribes multiple genes from common promoters, then one would expect that the colinear relation of co-transcribed genes would be preserved through evolution. Comparative genomic analysis of gene organization between *C. elegans and C. briggsae* – whose most recent common ancestor existed ∼100 Mya (Stein et al. 2003)– revealed that despite significant sequence divergence, ∼50% of all genes are arranged into blocks of perfect synteny where the exact orientation and positioning of orthologous genes are conserved (Vergara and Chen 2010). Because the arrangement of adjacent genes may be maintained if they share *cis*-acting regulatory sequences (i.e. a promoter/enhancer) (Quintero-Cadena and Sternberg 2016) it is not surprising that genes within the same operon (and share a common promoter) are typically maintained in the same syntenic blocks (Vergara and Chen 2010); yet such operons account for less than 1/3 of all syntenic genes in *C. elegans*. Considering the extent of contiguous, intergenic Pol II transcription, we investigated whether genes with the same transcription unit are likely to be maintained in the same syntenic region. Excluding genes within annotated operons (Allen et al. 2011), we found that 49% of colinear adjacent genes transcribed as part of the same polycistronic (>2) transcription unit are retained within the same syntenic block—this degree of conservation is significantly greater than the genome average of 34%.

The apparent relationship between transcribed genomic units and synteny of colinear genes prompted us to examine whether the conservation of the orientation and positioning of orthologous genes could be explained by transcription. While such analysis is confounded by the phylogenetic distance *C. elegans and C. briggsae*, and our limited understanding of all possible transcription patterns that can occur in *C. elegans*, we found repeated examples where one gene is apparently linked to a neighboring gene by transcription in *cis*. Typically, this included polycistronic transcription units, but also bidirectional promoters and many cases of convergent genes transcribed in antisense (Figure 6A, S4). We considered the possibility that such relationships may account for the retention of perfect synteny and chose to investigate this by collapsing overlapping transcription units on both strands into intervals that denote whether a genomic region is connected by continuous overlapping transcription (“transcribed region”, Figure 6A). We subsequently asked whether each gene in the genome is covered by a transcribed region; whether that gene was part of a syntenic block and if the gene was part of an operon. Figure 6B shows that 2/3 of all genes within syntenic blocks are found within multi-gene transcribed regions, a result not expected by chance (p<1_^10-10_, hypergeometric test). This group includes ∼80% of genes within annotated operons as well as >4000 other genes. Finally, we considered the possibility that the number of genes within a transcribed region should relate to the likelihood of those genes being retained in a syntenic block. As anticipated, regions containing a single transcribed gene showed modest enrichment above the genome average for being in a syntenic relationship (Figure 6C), but genes within multi-gene transcribed regions are increasingly more likely to have retained synteny with a nearby gene. Thus, it appears that adjacent genes are frequently linked by transcription and this association may underlie why the positioning and orientation of neighboring genes is preserved.

**Figure 6:**
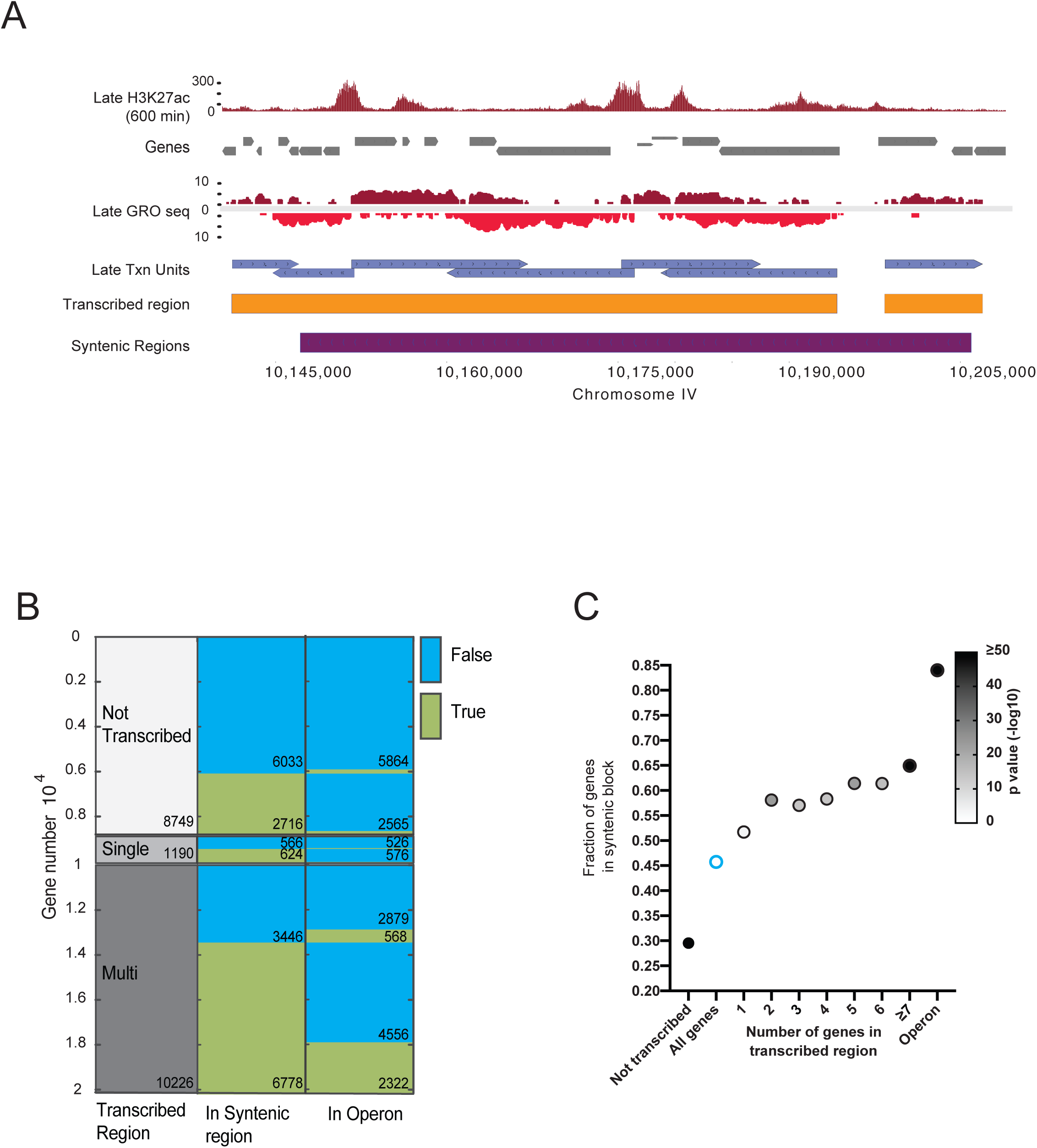
Overlapping transcribed regions define syntenic blocks between *C. elegans* and *C. briggsae*. a) Genome browser image that highlights nascent RNA transcription in late embryos. Top, histone modification patters. Middle, nascent RNA mapped by Gro-seq. Reads mapping to the Watson strand and Crick strands are shown above and below the center lines, respectively. Mapped transcription units are shown as blue boxes, the direction of the arrow indicates the strand on which the nascent RNA was mapped. Overlapping transcription units were collapsed into transcribed regions denoted as orange box. Regions of perfect gene synteny are shown as a purple box. b) Analysis of gene transcription with respect to operons and gene synteny. Left column, all annotated genes we classified as being in a transcribed region or not transcribed. Transcribed genes were further separated into those being part of single gene (single) or a multi gene (multi) transcribed region. Middle column, each gene was then classified as being in a syntenic block (green, true) or not (blue, false). Right column, each gene was then classified as being in an operon (green, true) or not (blue, false). Gene identity is maintained in rows that span the three columns. c) Multi gene transcribed regions are preferentially retained in syntenic blocks. Each gene in the genome was classified as being not transcribed, transcribed and part of a transcribed region, or part of an operon. Transcribed regions were then segregated according to the number of genes contained within the region (indicated on X-axis) and the fraction of genes within that transcribed regions that are also part of syntenic block (Y-axis). The fraction of all genes (not included in operon) that are classified as syntenic is shown as a blue oval. The fraction of genes within operons that are also syntenic are shown on the right (operon). Greyscale indicates p value calculated by hypergeometric distribution.

## DISCUSSION

### Transcriptional plasticity during *C. elegans* embryogenesis

*C. elegans* has acquired an exceptionally diversified RNA landscape with the functional capacity to bind, amplify, and degrade endogenous as well as exogenous transcripts through multiple RNA interference pathways (Billi et al. 2014). Recent small RNA profiling studies have indicated that essentially all transcripts are surveilled by the RNAi machinery (Seth et al. 2018; Shen et al. 2018). With so many layers of post-transcriptional regulation that are separated in developmental space and time, characterizing *C. elegans* RNA transcription simply by mRNA abundance is misleading. We reasoned that integrating independent maps of accessible chromatin, nascent RNA, and mature mRNA from developmentally synchronized embryos might reveal insights into the regulatory context to the *C. elegans* transcriptional regulation.

To help define the gene regulatory landscape of *C. elegans* embryogenesis, we purified chromatin from developmentally synchronized embryos to construct maps of engaged RNA polymerase II (Pol II) transcription at dramatically distinct morphological stages. We found that although embryos display diverse mRNA expression patterns as they develop (Packer et al. 2019a; Warner et al. 2019), chromatin states commonly associated with Pol II transcription initiation are broadly invariant and remain enriched at consistent genomic loci. These chromatin domains serve also sites for DNA replication initiation (Pourkarimi et al. 2016; Rodriguez-Martinez et al. 2017), centromere formation (Steiner and Henikoff 2014), and appear configured towards germ line gene expression programs, where mRNAs tend to encode essential, broadly-expressed proteins required for growth and often are transcribed from *C. elegans* operons (Reinke and Cutter 2009).

Using long-read nascent RNA sequencing, we were able to profile Pol II transcription prior to SL trans-splicing, revealing the extent and complexity of genome-wide transcription. While our data is in excellent agreement with previous reports that map nascent RNA, our ability to resolve early and late embryos has illuminated two key aspects of transcription: First, that Pol II can regularly transcribe clusters of genes in polycistronic transcription units outside of annotated operons; second, the constraint of initiating from developmentally conserved open chromatin domains requires Pol II to transcribe early expressed genes associated with growth and cell division, even in late embryos which are composed of non-replicating, differentiated or quiescent cells.

### Operons, pervasive transcription, and post-transcriptional mRNA control

*C. elegans* operons have been identified on the basis of SL2 containing mRNA transcripts and short intergenic (intercistronic) distances (Spieth et al. 1993; Blumenthal et al. 2002), yet the complete regulatory context of operons remains unclear. While the SL1 leader is typically assumed to be specific for genes at the 5’ end of the operon and the SL2 leader for those at the 3’ end, data suggest that trans-splicing is highly flexible— in certain contexts, leader sequences can substitute for each other (Spieth et al. 1993) or be excluded from the mRNA completely (Graber et al. 2007). Furthermore, SL-trans splicing is not restricted to genes in operons, as the majority of all mRNA transcripts (∼70%) have their 5’ UTR (i.e. outron) replaced by SL trans-splicing (Allen et al. 2011). Based on mRNA expression patterns from growth-related genes, ∼80% of which are in operons, it has been hypothesized that operons facilitate recovery from growth arrested states (e.g. dauer), as well as rapid transcription of mRNAs that support germ cell proliferation and oocyte production in reproducing populations of worms (Zaslaver et al. 2011). Unlike bacterial operons, *C. elegans* operons do not require mRNAs in the same transcription unit to be coexpressed, leading to the idea that operons have accumulated genes which can be controlled after transcription initiation (Blumenthal 2012). This model also proposed that genes in operons would be transcribed promiscuously from their promoters because the mRNAs would be regulated at the post-transcriptional or translational level, a phenomenon which we appear to detect as pervasive transcription of growth-related, genes from stable open chromatin domains (SACs).

Use of bulk nuclear assays such as Gro-seq and ChIP-seq do not allow us to ascribe specific chromatin states or nascent transcripts to individual cells or lineages of origin. One possibility is that pervasive transcription from consistently accessible chromatin domains is restricted to the minority of somatic cells (e.g. hypodermal, muscle, neuronal) which proliferate after hatching (Sulston and Horvitz 1977). However, the select inactivation of highly-transcribed early gene promoters would lead to an expected decrease in nascent RNA levels (Tome et al. 2018)—instead, we find consistent nascent RNA transcription, a readout shown to positively correlate with chromatin accessibility at multiple stages of *C. elegans* development (Daugherty et al. 2017).

Although orthologous RNA decay pathways controlling nascent and mature RNA transcript stability are largely undefined, operons contain more than 75% of *C. elegans* genes involved in mRNA degradation pathways (Blumenthal and Gleason 2003), suggesting that pervasive RNA transcription may be self-regulated by precise control of post-transcriptional RNA stability (Blumenthal 2012). We suspect that an RNA surveillance capacity encoded by endogenous RNAi pathways (Minkina and Hunter 2018; Seth et al. 2018) or microRNAs (Jan et al. 2011) might target the selective degradation of germline-expressed growth transcripts in embryonic tissues which do not resume proliferation after hatching (Almeida et al. 2019). Informed by stable transmission of an identical set of accessible chromatin sites, we are further interested in examining how pervasive RNA transcription in *C. elegans* might have evolved with other genome regulatory pathways, such as DNA replication, meiosis, and small RNA pathways (Claycomb et al. 2009; Gu et al. 2009; Gu et al. 2012; Pourkarimi et al. 2016). Particularly in an organism with a highly dynamic RNA regulatory landscape, the steady-state pool of pervasively transcribed RNA may supply endogenous siRNA (endo-siRNA) pathways with trigger sequences to stimulate tissue-specific mRNA surveillance across developmental stages (Pak et al. 2012; Newman et al. 2018). Alternatively, promiscuous transcription from open promoters has been hypothesized to support the biogenesis of small regulatory RNAs, such as miRNAs or piRNAs, which are known to guide the Argonaute class of RNA-binding proteins specifically to germline mRNAs (Kapranov et al. 2007; Gu et al. 2012; Shen et al. 2018).

### Chromatin maps, hyperaccessible regions, and enhancers

*C. elegans* clearly establishes new promoters and new sites of active histone modification as development proceeds (Daugherty et al. 2017; Janes et al. 2018), which we detect as the late appearance of H3K27ac and nascent RNA at annotated DAC regions (Figure 2a, Supplemental Table 2). The activation of these in the late embryo is coincident with the expected transcription of lineage-specific mRNAs (Levin et al. 2016) associated with terminal, differentiated cell fates (Boeck et al. 2016; Packer et al. 2019a). We detect highly similar patterns of intergenic Pol II transcription and histone modifications previously ascribed to tissue-specific developmental enhancers (Dupuy et al. 2004; Chen et al. 2013; Daugherty et al. 2017). This suggests that collective efforts to characterize the gene regulatory structure of *C. elegans* are capturing similar elements.

### Gene control without insulators

TADs are a common metazoan chromatin structure associated with gene looping interactions mediated by enhancers (Harmston et al. 2017)— although thought to be universal to metazoan gene regulation, many nematodes appear to have lost several components deemed to be functionally critical for gene regulation by chromatin looping, including chromatin insulators such as CTCF (Heger et al. 2009; Dekker and Heard 2015). Functional evidence that *C. elegans* enhancers can stimulate target gene transcription in a position and orientation independent manner is sparse. While identifying enhancers by assaying expression of heterologous reporters does permit efficient screening of *cis-*regulatory sequences (Okkema et al. 1993; Dupuy et al. 2004; Dupuy et al. 2007), such methods have caveats: first, enhancers are identified based on detection of reporter protein, which can be sensitive to gene copy number, RNA transcript stability, and translational efficiency; second, reporter assays lack measurements of Pol II transcription and so cannot readily define which step a putative enhancer might regulate gene expression. More current enhancer annotations in *C. elegans* are often based on a preferred combination of genomic readouts: histone modifications (histone acetylation, H3K4me), chromatin accessibility, and RNAs associated with Pol II initiation (e.g. promoter-associated RNAs, eRNAs) (Chen et al. 2013; Daugherty et al. 2017; Ho et al. 2017). But, given the questionable distinction between enhancers and promoters (Henriques et al. 2018), the understanding of enhancer function in *C. elegans* is incomplete (Barriere and Ruvinsky 2014; Janes et al. 2018). We suspect that gene regulation in *C. elegans* is generally not reliant on chromatin insulation or classical gene enhancers that can function irrespective of orientation and position that are used in other eukaryotes. Rather gene activity may be fundamentally regulated by directional *cis*-acting elements (i.e. promoters) (Henriques et al. 2018; Mikhaylichenko et al. 2018) and post-transcriptional mechanisms that control mRNA abundance.

Ancient nematode genomes, such as *Trichinella spiralis*, do encode CTCF, leading to a proposal that the CTCF gene was lost during the emergence of the derived, crown clade of nematodes, to which *C. elegans* and *C. briggsae* belong (Heger et al. 2009). Consistent with the loss of CTCF, the *C. elegans* genome does not appear to follow conserved nature of Hox gene evolution, which exhibits a spatiotemporal expression pattern largely maintained through chromatin insulation and enhancer looping (Ruvkun and Hobert 1998; Aboobaker and Blaxter 2003). Without the ability to stabilize chromatin looping domains through insulation, the *cis-*regulatory sequences controlling *C. elegans* Hox gene expression may have become influenced by neighboring patterns of transcription and acquired new regulatory functions (Cowing and Kenyon 1996; Aboobaker and Blaxter 2003).

The ability to trans-splice has likely relaxed the requirement for each gene to maintain its own its own 5’ *cis*-regulatory module (Blumenthal et al. 2015) leading to the linking of transcription patterns through colinear gene arrays (Longman et al. 2000; Huang et al. 2001; Blumenthal 2012). Our annotation of the nascent RNA landscape prior to trans-splicing suggests the existence of many transcription units (n=>1500) that behave similar to operons, the extent of intergenic transcription suggests Pol II elongation and initiation dynamics at distinct embryonic or larval stages might be influenced by interactions between the nascent transcriptome and RNA regulatory pathways, i.e. alternative/trans-splicing (Longman et al. 2000; Zahler 2005) and nuclear RNAi (Guang et al. 2010; Wedeles et al. 2013). Providing that an open reading frame can be transcribed, control of mRNA levels – and gene expression – would be achieved through the development of an extensive toolkit to modulate RNA post-transcriptionally. A notable feature of *C. elegans* biology is the extent of post-transcriptional control, especially focused towards the 3’ UTR sequences of germline-expressed transcripts (Merritt et al. 2008; Flora et al. 2018; Newman et al. 2018; Seth et al. 2018). 3’ UTR sequences may be directing SL trans-splicing proteins, RNA binding proteins, or Argonauts to interact with nascent pre-mRNA transcripts to guide appropriate post-transcriptional fates (Jager et al. 2007; Mangone et al. 2010; Newman et al. 2018; Shen et al. 2018).

### Transcription regulation has shaped the genome

Orthologous genes often remain linked on chromosomes in clusters for long evolutionary periods, however the mechanisms that establish and maintain genes in synteny remain less clear (Caron et al. 2001; Lercher et al. 2002; Vavouri et al. 2007). Analysis of gene order conservation across Eukaryotic genomes indicates that highly conserved gene pairs are retained in proximity due to shared transcriptional regulation (Davila Lopez et al. 2010). Larger syntenic blocks in vertebrate (Kikuta et al. 2007) and insect (Engstrom et al. 2007) genomes are often flanked by regulatory elements that function as enhancers of developmental gene expression, thus creating a selective pressure to maintain the chromosomal positioning of these sequence. However, the developmental constraint to maintain precise gene arrangement is presumably lessened by enhancers’ ability to activate target genes at varying distances and orientation.

Despite having diverged ∼100 million years ago, *C. elegans* and *C. briggsae* have retained a remarkable of number of orthologous genes in regions that maintain identical order and strandedness i.e. perfect synteny. Such precise conservation indicates that *C. elegans* and *C. briggsae* are intolerant of genome rearrangements that alter the orientation of neighboring genes. We suggest that distinct features of *C. elegans* gene regulatory logic might explain the maintenance of gene order (Figure 7). First, the lack insulator proteins and of TAD-like structures in *C. elegans* indicate that gene regulation is generally not reliant on chromatin insulation or classical gene enhancers that can function irrespective of orientation and position. Second, widespread utilization of SL trans splicing permits multiple, adjacent genes to be transcribed under the control of single promoter (Allen et al. 2011); including genes in operons (Zorio et al. 1994), and thousands of other colinear gene pairs that that are apparently transcribed together. Third, extensive use of post-transcriptional control mechanisms alleviates the need to regulate the expression of each gene by transcriptional activation or repression. Fourth, widespread use of bidirectional initiation from divergent promoters (Janes et al. 2018) would permit transcription of two adjacent clusters of genes from a single regulatory site. Finally, antisense Pol II transcription that converges into overlapping 3’ UTRs (Merritt et al. 2008; Mangone et al. 2010; Jan et al. 2011) links adjacent genes and their regulatory elements together. Thus, *C. elegans* is likely to be particularly sensitive to the maintenance of gene position relative to *cis*-acting sequences. It is noteworthy that trypanosomes, which make extensive use of polycistronic transcription and post-transcriptional mRNA control, also exhibit a striking conservation of gene order over vast evolutionary timescales (∼400-600 Myr) (Ghedin et al. 2004).

**Figure 7:**
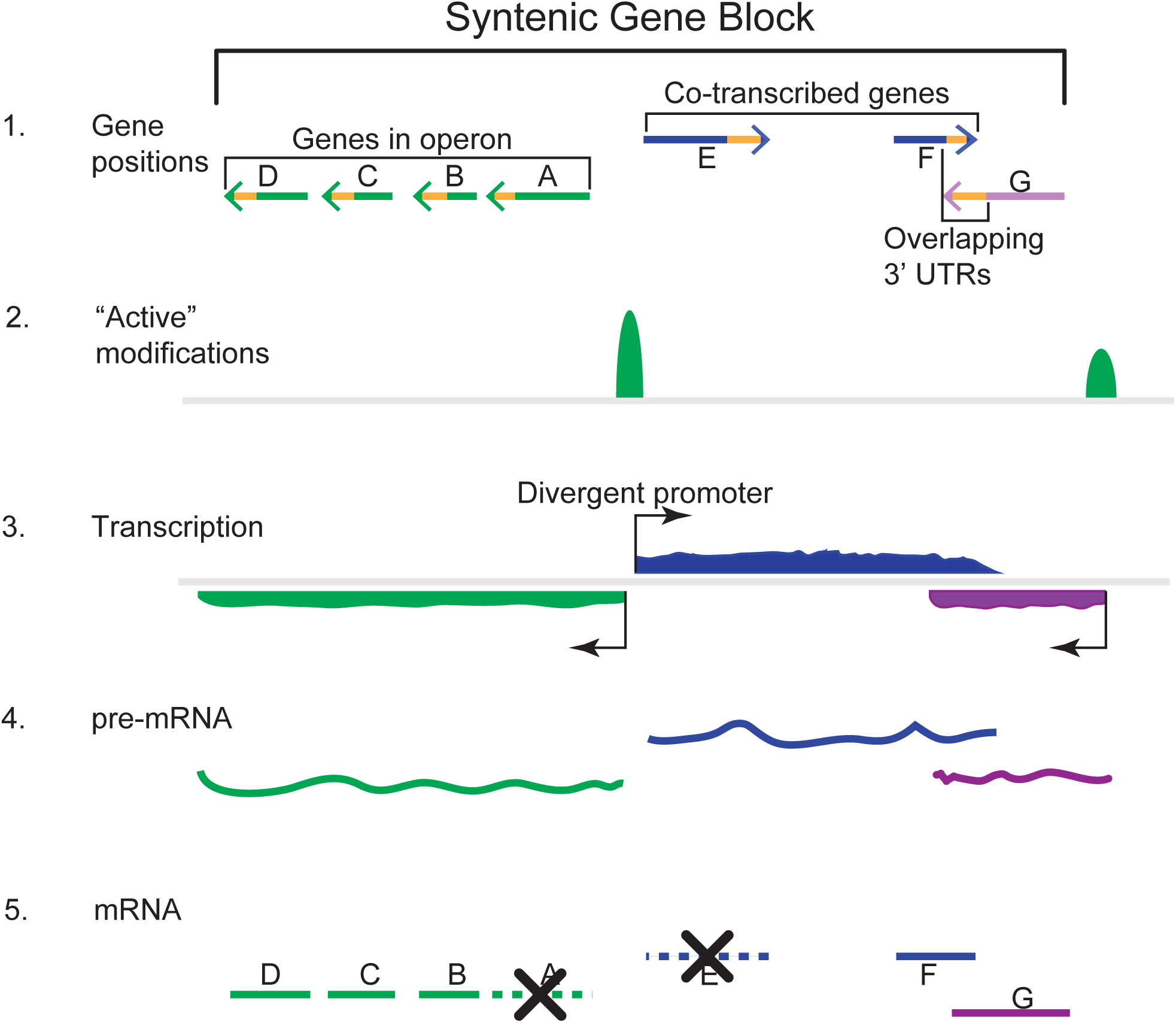
Overlapping transcribed regions define syntenic blocks. Model to explain how overlapping transcription and shared *cis*-regulatory sequences can link adjacent genes in syntenic blocks: 1) hypothetical gene arrangement showing genes in operon (green), co-linear genes separated by large intergenic space (blue), single gene whose 3’UTR overlaps with an adjacent gene (purple). Regulatory elements such as 3’ UTRs shown as yellow. 2) active histone modifications i.e. H3K4me2, H3K27ac, mark sites of transcription initiation. 3) transcription of multiple genes occurs from a limited number of promoters – creating pre-mRNA that may contain a number of coding sequences, 4-5) the pre-mRNA is processed by SL-trans splicing machinery to generate individual mRNA molecules; these are subject to RNA surveillance that degrades unwanted transcripts, donated by a dashed line and an X.

The gene regulatory system developed during *C. elegans* natural history has undoubtedly influenced the organizational structure of the genome. In particular, the absence of a system of chromatin insulation to compartmentalize Pol II transcription activity may have reduced the functional capacity of enhancers to regulate gene expression timing across broad chromosomal domains—such a deficiency may have led to an increased reliance on *cis*-regulatory elements within colinear transcription units to control gene expression through post-transcriptional RNA pathways. It will be of great interest to understand how RNA Pol II elongation and termination are controlled, considering the prevalence of shared promoters and co-transcription of neighboring genes.

## METHODS

### *C. elegans* maintenance and strains

All *C. elegans* strains were maintained at 20°C on NGM plates with OP50 *E. coli* strain, as previously described (Brenner, 1974). The early egg-laying mutant strain, CG21 *egl-30(tg26) I; him-5(e1490) V,* was used to harvest embryos for chromatin and RNA sequencing libraries (Pourkarimi et al. 2016).

### Bleaching and *in vitro* staging of synchronized embryos

Approximately 350,000 L1-stage *egl-30(tg26)* worms were grown to adulthood at 20°C on NGM plates until early embryos were detectable *ex utero*, when adults were washed from plates into 50 mL Falcon tubes with M9 buffer. Adults settled to the bottom of the tube by gravity and were washed a total of 5 times with fresh M9 until most *ex utero* embryos were washed from solution (Pourkarimi et al. 2016). Embryos were bleached with sodium hypochlorite solution, washed with M9 buffer, and resuspended in aliquots of egg buffer (Edgar et al. 1988). Aliquots were incubated on a rotating stand at 20°C until the desired developmental time was reached (Early: 60 minutes, Gastrula: 200 minutes, Late: 600 minutes).

Proper embryo synchrony and developmental staging were visually confirmed by microscopy during each time-course experiment—developmental timing was validated by comparing mRNA transcripts in staged embryo populations to expected gene expression patterns from single-embryo studies (see Supplemental Figures) (Hashimony et al. 2015).

### ChIP-seq

Staged embryos designated for ChIP-seq analysis were cross-linked with a 2% formadehyde solution (diluted in 500 mL M9 buffer) for 30 minutes, followed by the addition of 125 mM glycine to quench the reaction, and two final washes in PBS + protease inhibitors. Extracts were prepared from synchronized *egl-30(tg26)* embryos by standard methods described previously (Reichsteiner et al. 2010, Ercan et al. 2007). Approximately 2 mg of extract from each embryo stage was immunoprecipitated with 2 ug of commercially available antibodies previously cited in *C. elegans* chromatin studies: H3K27ac (Abcam: ab4729) (Liu et al. 2011) and H3K4me2 (Abcam: ab7766) (Xiao et al. 2011). Extracted DNA was prepared into ChIP-seq libraries using the NEBNext Ultra DNA library preparation kit for Illumina (New England Biolabs) and was sequenced on the HiSeq 2500 platform.

### Genomics: Chip-seq analysis

Throughout the mapping of ChIP-seq reads to the genome, standard sequencing quality control steps were conducted. Individual sequencing reads in early embryo, gastrula, and late embryo replicates were annotated into peak calls using MACS (v2.1) (--broad peak call) (Zhang et al. 2008). We considered high-confidence H3K27ac and H3K4me2 peaks to be those which were: consistent across replicate time points, detected by sonication and MNase assays (Supplemental Figure 1), or supported by previously published *C. elegans* chromatin annotations (Gerstein et al. 2010, Daugherty et al. 2017, Ho et al. 2017).

The stability of chromatin states across embryogenesis was determined using the bedtools— high-confidence histone modification peaks detected in gastrula and late embryos (overlap = 0.25) were classified as stable active chromatin (SAC) domains (Supplemental Table), while peaks which only appear in late embryos were characterized as dynamic active chromatin (DAC) domains (Supplemental Table).

### Global run-on sequencing with Oxford Nanopore MinION

Staged embryos designated for Gro-seq analysis were prepared as described previously in Kruesi et al. 2013, with minor alterations. First, we did not fragment nascent BrUTP-labeled RNA transcripts with sodium hydroxide, electing instead to keep full-length, nascent RNA molecules intact. Second, we used the Ribo-Zero Gold rRNA Removal Kit (Illumina) to deplete rRNA transcripts from immunoprecipitated BrUTP-labeled nascent RNA in an effort to minimize noise and increase transcriptome coverage in our Gro-seq libraries.

Sequencing libraries were prepared from BrUTP-labeled RNA using the PCR-cDNA sequencing kit from Oxford Nanopore Technologies (cat: SQK-PCS109). Libraries were prepared using ∼100 ng of input RNA according to the manufacturer’s specifications. Briefly, full-length nascent RNAs are polyadenylated *in vitro* to initiate annealing of 3’ poly-T oligo and reverse transcription by SuperScript IV Reverse Transcriptase (cat: 118090010). Double-stranded cDNAs were size-selected to a range of 300-1000 bp using SPRI beads (AmpureXP) and were amplified with low-cycle PCR prior to sequencing. Following limited PCR amplification, cDNA libraries were primed with rapid sequencing adapters and loaded onto the MinION flow cell. We sequenced independent, replicate cDNA libraries from gastrula and late stage embryos—we later merged reads from gastrula and late stage replicate libraries the Gro-seq profile at each developmental time point consists of ∼15 million total reads.

### Genomics: Alignment of Gro-seq reads

Nanopore reads were converted into FastQ using Guppy (Oxford Nanopore). Strand information of the sequenced cDNA was deduced by identifying known adapter sequences at each end of the sequence read using Last (Genome Res. 2011 21(3):487-93). Adapter indexes were created for: ONT_VNP ACTTGCCTGTCGCTCTATCTTCTTTTT and ONT_SSP TTTCTGTTGGTGCTGATATTGCTGGG using the lastdb function. The indexes were than matched to the fastQ reads using: lastal -Q 1 -P10. Reads with duplicated adapters and reads with adapters not within 80nt of end of sequence were discarded. The small fraction of reads matching splice leader sequences: CE_SL1, GGTTTAATTACCCAAGTTTGAG, CE_SL2 GGTTTTAACCCAGTTACTCAAG were also identified using Last and discarded. Strand-specific fastQ files were then mapped to the Ce10 genome using Minimap2 with standard parameters for nanopore data (-ax map-ont). Read abundance for each gene was calculated using Bedtools with the WS235 reference genome.

### Annotation of transcription units using gro-HMM

Continuous stretches of Gro-seq signal were annotated using gro-HMM as previously described (Chae et al. 2015). Briefly, nascent RNA sequencing files (.bam format) were aligned to the WS235 genome and converted to genomic ranges objects in the R statistical environment. Transcripts are detected using a two-state hidden Markov model, based on calculating transitions from “transcribed” to “non-transcribed” genomic regions—based on author recommendations, we adjusted model parameters to account for the smaller genome size and clustered gene spacing found in *C. elegans* (LtProb = −75, UTS = 5) (Chae et al. 2015).

To limit the contribution of transcriptional noise into our annotations of Pol II transcription, we required each gro-HMM unit to overlap the coding sequencing of annotated WS235 genes by at least 25%. Genes were assigned into a transcription unit if 50% of their coding sequence was covered by a transcription unit.

---

**Figure S1.**
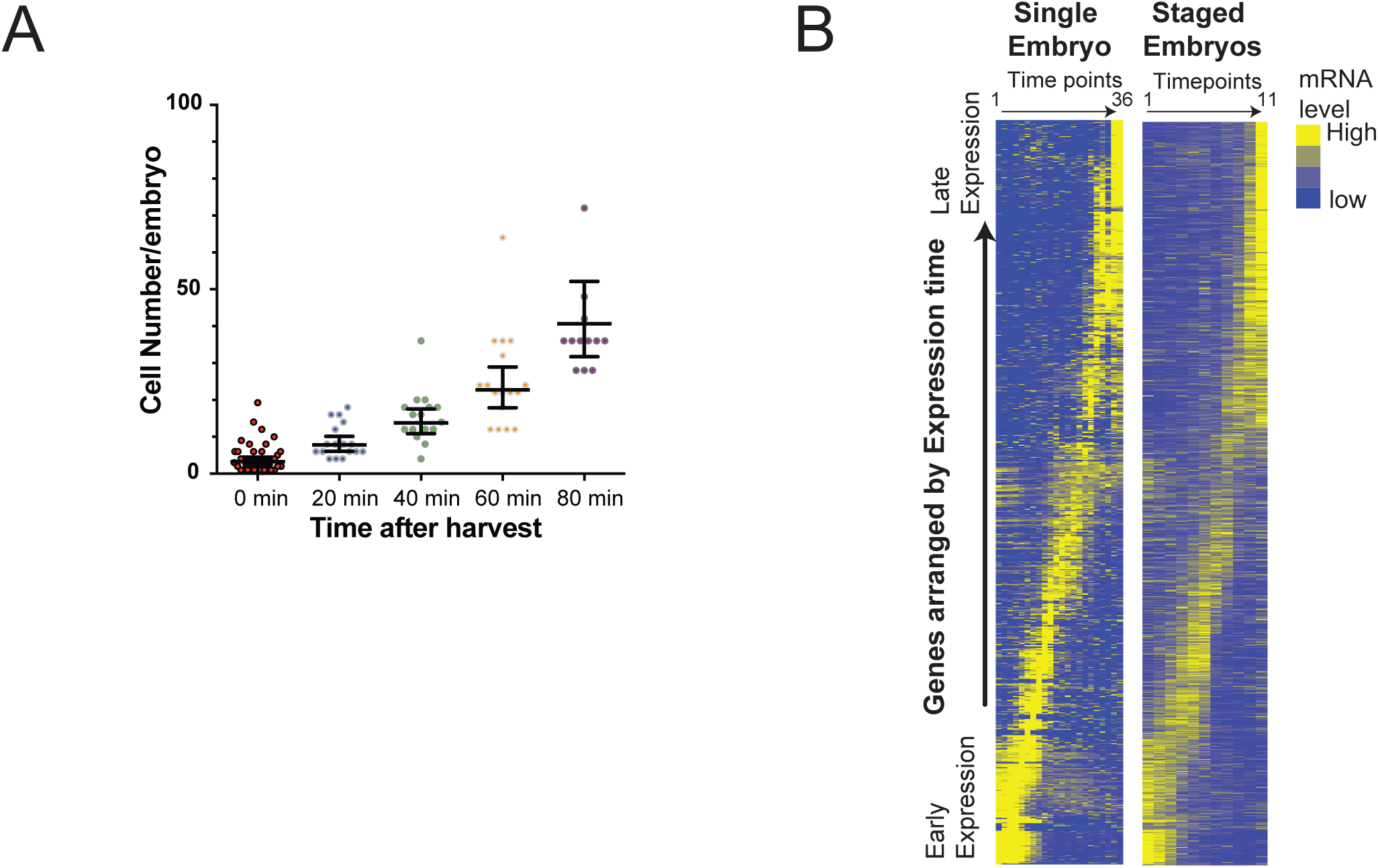
A, Embryo cohorts from early egg-laying *egl-30(tg26)* captures developmental timing of *C. elegans* embryogenesis. Cell number was determined by DAPI staining. After harvest, (0 min) embryos were allowed to develop for various times (x-axis) at room temperature in standard buffer. Black bars represent geometric mean and 95% confidence intervals. B, Heatmaps showing transcript abundance in single embryo (Hashimony et al. 2015, left) and highly synchronized embryo populations (this study, right) through the proliferative stage (first 420 minutes) of embryo development. Genes are ranked according to maximal expression time in the single embryo time course. High abundance transcripts in the earliest time points that later decay are maternally deposited. All genes are normalized such that the sum of expression through the time course =1.

**Figure S2.**
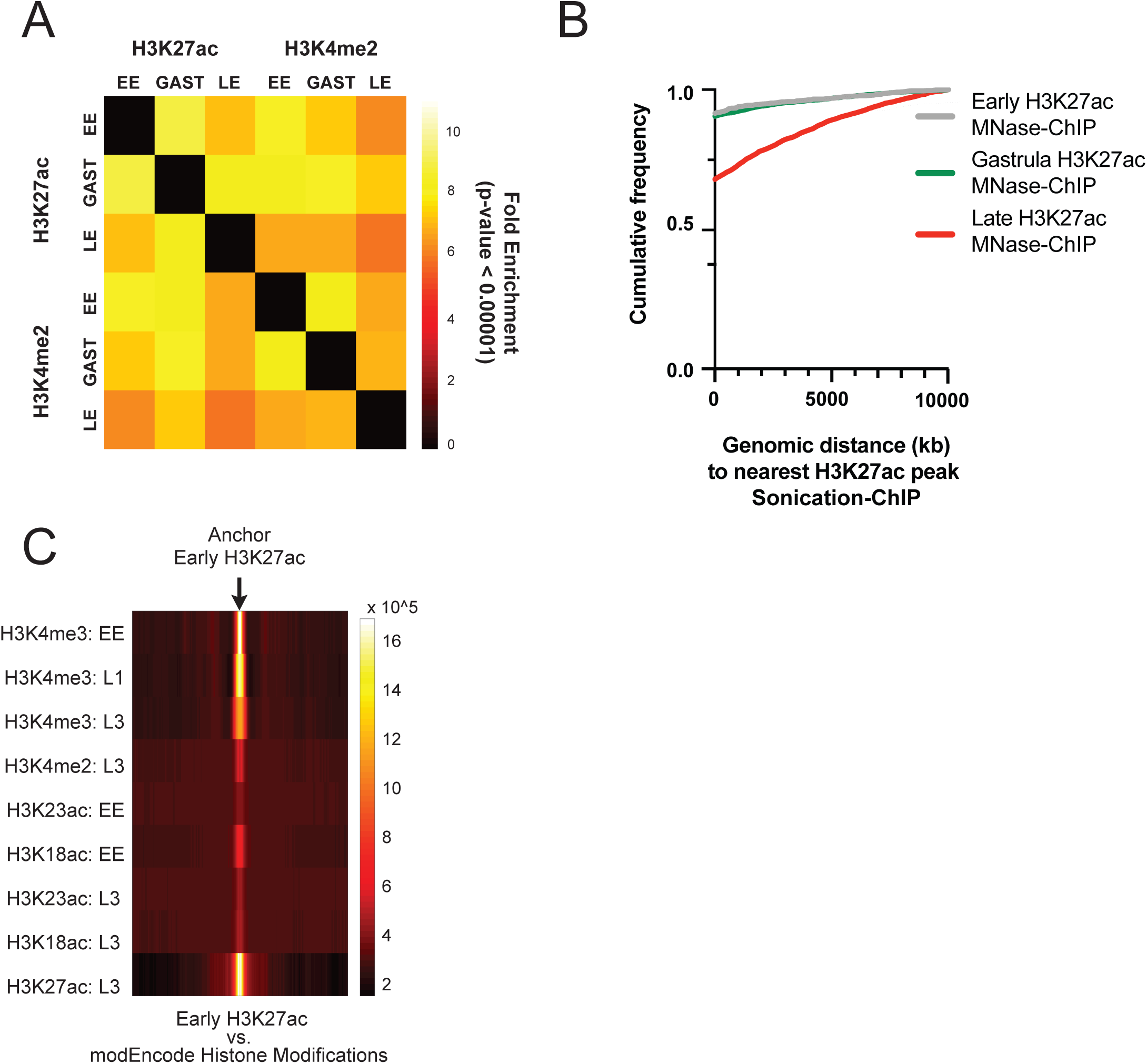
A, Consistency in *C. elegans* active histone modification maps. Pairwise comparison of the overlap between histone modifications mapped in the embryo. Heatmap indicates fold enrichment of overlapping regions of histone modification. All indicated overlaps are not expected by chance: p-value < 0.00001, hypergeometric distribution. B, The distance between significantly enriched genomic intervals mapped by MNase-ChIP and regions mapped by sonication-ChIP is shown. Colored lines denote the developmental stage at which MNase-ChIP against H3K4me2 was performed. The distance between the midpoint of all enriched regions (defined by MACS v2.1) is calculated, binned and summed. C, Comparison of regions with enriched histone modifications in embryonic and larval stages defined by modEncode. Regions enriched for H3K27ac in the early embryo are used as the anchor and the abundance of ChIP signal for the indicated modifications and developmental stage is mapped for a 50KB region centered on the anchor. Color scale denotes the abundance of mapped signal from the indicated histone modifications on left.

**Figure S3.**
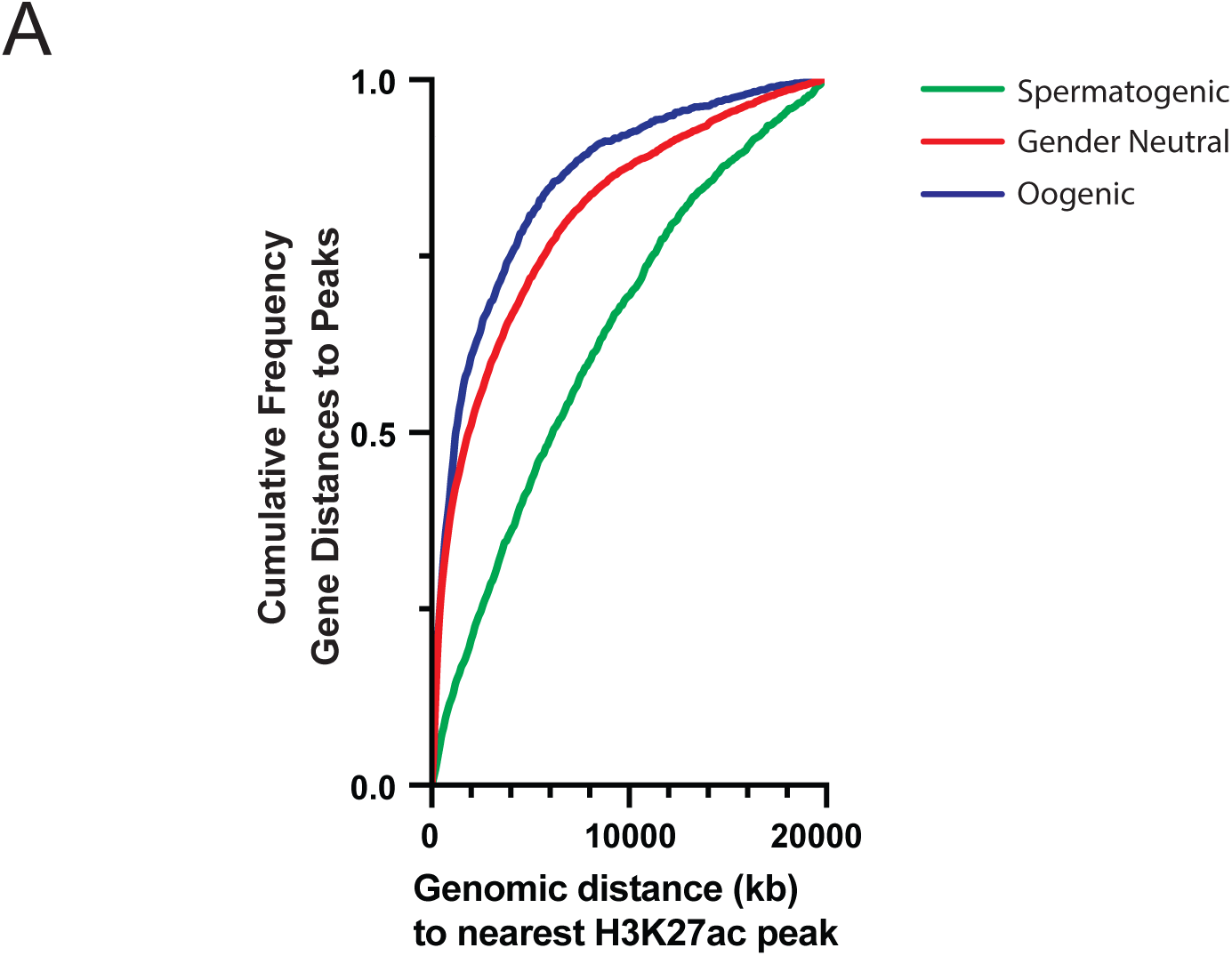
Association of germline-expressed genes with earliest embryonic histone modifications. The distance between genes predominantly expressed in the male (spermatogenic, green) female (oogenic, blue) or male and female (gender neutral, red) and regions enriched for H3K27ac in the early embryo is shown (nearly identical results for H3K4me2, data not shown). The distance between the start of the first exon and the midpoint of the nearest enriched region or H3K27ac (defined by MACS v2.1) is calculated, binned and summed. Germline expression data is from Ortiz et al. 2014.

**Figure S4.**
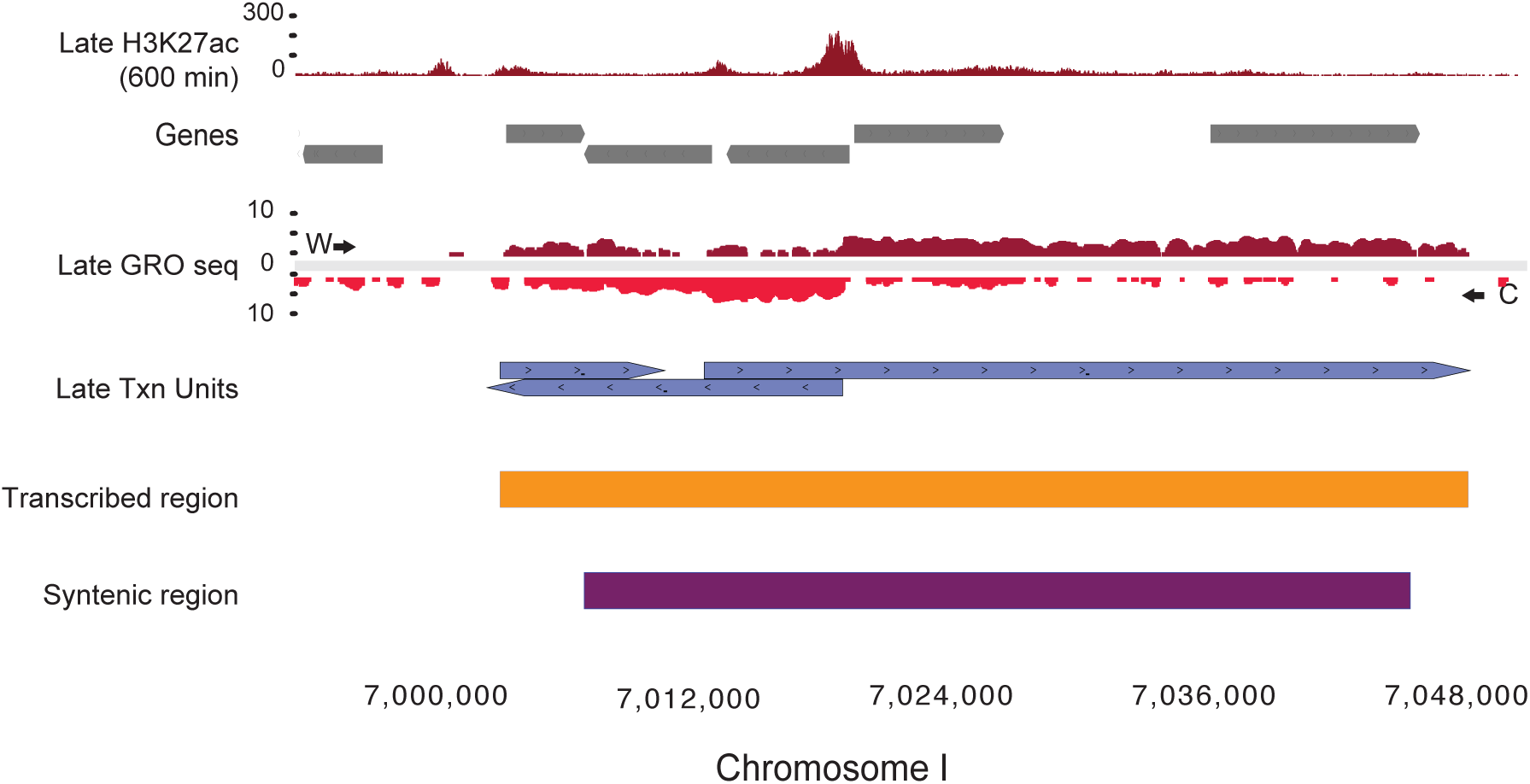
Genome browser image that highlights nascent RNA transcription late embryos. Top, histone modification patterns. Middle, nascent RNA mapped by Gro-seq. Reads mapping to the Watson strand or Crick strands are shown above and below the center lines, respectively. Mapped transcription units are shown as blue boxes, the direction of the arrow indicates the strand on which the nascent RNA was mapped. Overlapping transcription units were collapsed into transcribed regions denoted as orange box. Regions of perfect gene synteny are shown as a purple box.

**Supplemental Table 3a.** Gene Ontology: DAC-associated genes

| Gene Ontology Category | p-value |
| --- | --- |
| actomyosin structure organization (GO: 0031032) | **6.29 E-07** |
| positive regulation of axonogenesis (GO: 0050772) | **5.34 E-06** |
| cilium organization (GO: 0044782) | **8.86 E-08** |
| chemotaxis (GO: 0006935) | **5.64 E-07** |
| locomotion (GO: 0040011) | **3.75E-06** |

**Supplemental Table 3b.** Gene Ontology: SAC-associated genes

| Gene Ontology Category | p-value |
| --- | --- |
| DNA replication initiation (GO:0006270) | **6.95E-04** |
| mRNA 3'-end processing (GO:0031124) | **3.39E-04** |
| mitotic sister chromatid segregation (GO:0000070) | **3.90E-08** |
| nuclear envelope organization (GO:0006998) | **9.31E-05** |
| macroautophagy (GO:0016236) | **1.29E-10** |

